# SP1 tethers NR2F2 to drive atrial cardiomyocyte identity

**DOI:** 10.64898/2026.10.19.760167

**Authors:** Payton N. Gergen, Alexander P. Clark, Jeffrey J. Saucerman

## Abstract

Human heart development is guided by complex gene regulatory networks (GRNs), which are difficult to unravel and differ from that in animal models. In this study, we use a systematic approach to identify gene interactions driving human fetal atrial cardiomyocyte identity. We inferred GRNs for atrial and ventricular cardiomyocytes from human fetal single-cell RNA-seq and ATAC-seq data, predicting transcription factors that differentially regulate atrial vs. ventricular cardiomyocyte identity. Comparison of GRNs constrained by ATAC-seq vs. promoter annotations revealed that during human heart development, the canonical transcription factor for atrial specification, NR2F2, drives atrial genes not directly but by interacting with SP1. The predicted NR2F2-SP1 relationship at atrial gene promoters was supported with AlphaFold predictions of biophysical tethering and comparison with previous ChIP-seq data. These predictions were experimentally validated in human iPSC-derived atrial cardiomyocyte differentiation, which was suppressed by SP1 knockdown. Further, proximity ligation experiments validated the physical interaction of NR2F2 and SP1 in the nucleus, coincident with atrial identity. This study reveals SP1 as a transcription factor necessary for atrial cardiomyocyte identity in human development, helping to coordinate the effects of NR2F2. Furthermore, comparing GRNs based on current and potential epigenetic state offers a novel systematic approach to study interactions between transcription factors, which is key to understanding complex GRNs.

## Introduction

The heart is the first functional organ to form, elaborating multiple distinct cellular states and tissue morphologies. One critical bifurcation in heart development is the emergence of distinct atrial and ventricular subtypes. Atrial and ventricular cardiomyocytes have thousands of differentially expressed genes, leading to distinct structures and electrophysiological properties.^1–4^ These distinct atrial and ventricular cellular identities are regulated at the molecular level by an array of transcription factors (TFs) that bind at open chromatin in gene enhancers or promoters, forming complex gene regulatory networks (GRNs). Mutations in TFs such as pro-atrial (NR2F2^5,6^ and TBX5^7–9^) and pro-ventricular (HEY2^10–12^ and IRX4^13,14^) have been shown to induce an array of human congenital heart defects. However, due to the complexity of gene regulation, these studies have largely focused on the effects of individual TFs on target expression and cardiac phenotypes.

Single-cell RNA sequencing has enabled genome-wide analysis of cardiac gene expression during development in cultured cells,^9,15^ mice,^16,17^ and human fetal heart.^4,18^ While gene expression alone can identify correlated gene sets or infer trajectories, its combination with single-cell chromatin accessibility enables genome-wide inference of mechanistic GRNs.^17,19–22^ Here, we leverage such data to develop mechanistic GRNs for human fetal atrial and ventricular cardiomyocytes, with which we identify regulators of atrial cardiomyocyte identity.

Although targeted and genome-wide studies have largely examined transcription factors (TFs) independently, TFs from distinct structural families can cooperate to regulate gene expression in a DNA sequence-dependent manner.^23^ TF-TF screens have identified thousands of TF interactions at DNA,^24^ and some TF-TF interactions have been shown to regulate cell fate or subtype specification.^25,26^ Systematic approaches are needed to discover such TFs that cooperate to modulate the identity of cardiomyocytes in the developing heart.

In this study, we leveraged single-cell transcriptomics and chromatin accessibility of the human fetal heart to map TFs and their interactions that regulate chamber-specific cell identity. Comparison of gene regulatory networks constrained by promoter annotations vs. chromatin accessibility predicted that NR2F2, the only gene characterized to be necessary and sufficient for specifying atrial identity,^6,7^ does so through cooperating with SP1. Using AlphaFold3, we predicted a biophysical interaction of NR2F2 and SP1 important for tethering of NR2F2 at atrial gene promoters. Both NR2F2-SP1 nuclear interaction and the necessity of SP1 for atrial gene expression were experimentally validated in human iPSC-derived cardiomyocytes.

## Methods

### Inference of gene regulatory networks

We acquired pre-processed scRNA-seq data of human fetal heart samples obtained at post-conception weeks 6, 8, 12, and 19 from published datasets (GEO GSE181346) ^4,18,27,28^, from which we extracted time-pooled aCM data and time-pooled vCM data. Similarly, we extracted time-pooled aCM and time-pooled vCM snATAC-seq acquired from human fetal heart at post-conception weeks 6, 8, and 19^29^. Using these data, gene regulatory networks for human fetal aCMs and vCMs were inferred using CellOracle (v0.20.0)^20^ using parameters specified below.

#### ATAC-informed GRN construction

First, aCM and vCM scATAC-seq were pooled to construct a fetal cardiomyocyte base GRN, constraining potential TF-gene interactions to regions of open chromatin. Peaks were called separately for aCMs and vCMs with MACS2 and merged into a peak set (hg38) using ArchR. To construct the base GRN, CellOracle uses HOMER-derived TSS annotations^30^ to identify proximal regulatory DNA elements and Cicero^31^ to identify distal regulatory DNA elements in the accessible regions as indicated in the scATAC-seq. Peaks with a Cicero co-accessibility score >0.8 to a TSS peak were retained. CellOracle then scans the regulatory DNA sequences for TF binding motifs using gimmemotifs (gimme.vertebrate.v5.0.motifs, false positive rate = 0.02, motif score threshold = 10)^32^ to create a list of potential regulatory interactions. The base GRN was then used together with the TF-gene regularized regression from the aCM scRNA-seq to infer edge weights for the aCM GRN. Likewise, the base network was used with vCM scRNA-seq to infer edge weights for the vCM GRN. GRN inference was performed using default CellOracle parameters. The optimal number of nearest neighbors for KNN imputation was 66. The sensitivity for edge inference was alpha = 10. Each GRN was then filtered to the top 2,000 edges based on absolute edge weight out of >60,000 edges, with all filtered edges having p < 0.001.

#### Promoter-informed GRN construction

The promoter-informed GRN was based on the pre-built CellOracle human base GRN, which uses human genome hg38 with gimmemotifs^32^ (vertebrate motifs v5.0) to identify TF motifs in gene promoters. Unlike the ATAC-informed base GRN, the pre-built base GRN is not cell type-specific. Promoter-informed aCM and vCM GRNs were then inferred using the promoter-informed base GRN together with aCM or vCM scRNA-seq, respectively, and the same parameters as the ATAC-informed networks. The GRNs were each filtered to the top 2,000 strongest edges out of >40,000 total edges, with all filtered edges having p < 0.001.

#### Predicting drivers of atrial and ventricular identity

We determined the top 20 atrial and top 20 ventricular DEGs from the pooled aCM and vCM scRNA-seq by a Wilcoxon signed-rank test. For each source TF in the aCM GRN, we summed the weight of all edges from that TF to top aCM DEGs. We repeated this process with TFs targeting vCM DEGs in the aCM GRN, then did the same for the vCM GRN. These metrics provided a quantitative summary of the role of a TF in regulating both atrial and ventricular genes in aCMs and in regulating both atrial and ventricular genes in vCMs.

To rank the overall influence of each TF on atrial and ventricular genes in aCMs, an atrial-ventricular net identity score was calculated by subtracting the total weight of TF edges to atrial genes from the total weight of TF edges to ventricular genes. A more negative score indicates that the TF is pro-atrial (promotes atrial genes and/or represses ventricular genes), while a more positive score indicates that the TF is pro-ventricular (represses atrial genes and/or promoters ventricular genes). Scores nearer to 0 indicate little net effect on atrial vs. ventricular identity.

### Protein structure modeling

We used AlphaFold3^33^ to predict the structure and interaction of SP1 and NR2F2 with double-stranded DNA sequences at a PAM, TBX5, DKK3, NES, or CPNE5 promoters. These genes were selected from the top 20 differentially expressed atrial genes, were NR2F2 targets in the promoter-based aCM network, and were SP1 targets in the ATAC-based aCM network. We then used Find Individual Motif Occurrences (FIMO) to locate the top SP1 binding motifs. For the AlphaFold predictions, we input 50-base pair sequences centered at the strongest SP1 binding motif identified using FIMO and the reverse complement of the 50 bp strand. Controls included the predicted structure of NR2F2 alone, SP1 alone, NR2F2 and SP1, NR2F2 with atrial promoter DNA, and SP1 with atrial promoter DNA. We also predicted the structure of SP1 and NR2F2 with scrambled atrial promoter sequences and with random sequences of DNA with varying GC-content.

Of the five AlphaFold predictions for each set of inputs, we selected the predicted structure with the lowest fraction of disordered content for subsequent analysis. We used a pair interface-predicted template modeling (iPTM) score threshold of 0.1 to determine whether protein-protein or protein-DNA binding was predicted. We visualized AlphaFold predictions in PyMol,^34^ using predicted local distance difference test (pLDDT) score to identify regions of highest confidence. We used the PyMol visualizations to identify regions on SP1 where the protein interacted with DNA. We also calculated solvent-accessible surface area per molecule for SP1, NR2F2, and the DNA segment in PyMol.

### Analysis of ChIP-sequencing data

As an independent assessment of whether SP1 indeed binds at the accessible regions of promoters of atrial genes, we examined publicly available ChIP-sequencing data. We compared HCT116 SP1 (ENCSR000BSF) accessed through the ENCODE portal^35,36^ with the aCM ATAC-seq used in building the ATAC-informed aCM GRN. All datasets were aligned to hg38 and visualized in IGV^37^. Promoter labels were accessed from EPDnew^38^. We examined binding in the accessible regions of the promoters of atrial genes PAM, TBX5, DKK3, NES, and CPNE5. We called peaks using MACS2^39^.

### Cardiomyocyte experiments

#### Human iPSC-cardiomyocyte differentiation

GFP-tagged MYL2 WTC11cells (AICS-0060-027) were plated in Matrigel-coated 6-well plates until reaching >90% confluency. The iPSCs were then differentiated toward atrial cardiomyocytes using the STEMdiff™ Atrial Cardiomyocyte Differentiation Kit (STEMCELL Technologies #100-0215) according to manufacturer instructions until 20 days of differentiation.

#### Knockdown of SP1 in hiPSC-CMs

SP1 was knocked down using third-generation, self-inactivating lentiviral vectors that express shRNA (VectorBuilder; Scramble, VB250915-1419cvs with TagBFP2 in place of EGFP; SP1, VB250915-1419cvs with TagBFP2 in place of EGFP). In each vector, the human U6 promoter drives the shRNA. A human PGK promoter drives TagBFP2 fused to a puromycin resistance so transduced cells can be identified by blue fluorescence.

hiPSC-aCMs differentiated for >20 days were dissociated (STEMdiff Cardiomyocyte Dissociation Kit) and replated into (Matrigel-coated) 96-well plates at 20,000 cells/well. The next day, cells were transduced by overnight incubation with lentivirus in maintenance medium. Medium was replaced the next day and every 2–3 days after that. Cells were analyzed 7 days after transduction.

#### Immunofluorescence

hiPSC-CMs were fixed in 4% PFA for 20 minutes and washed with PBS. Cells were permeabilized in 0.2% TritonX-100 for 15 minutes at room temperature then incubated in 1% BSA for 20 minutes at room temperature to block nonspecific binding. Cells were then incubated in primary antibody solution for 90 minutes at room temperature. The primary antibodies used were 1:250 mouse monoclonal cardiac Troponin T (Thermo Fisher Scientific, MA5-12960), 1:250 goat polyclonal ANP (R&D Systems, AF3366), and 1:167 rabbit polyclonal MLC2V (Abcam, ab79935).

After washing with PBS and incubation in 5% donkey serum for 5 minutes at room temperature for blocking, cells were incubated in secondary antibody solution for 45 minutes at room temperature. Secondary antibody solution included 1:200 donkey anti-mouse IgG, Alexa Fluor^TM^ 488 (Thermo Fisher Scientific, A21202); 1:200 donkey anti-goat IgG, Alexa Fluor^TM^ 647 (Thermo Fisher Scientific, A21447); and 1:200 donkey anti-rabbit IgG, Alexa Fluor^TM^ 594 (Thermo Fisher Scientific, A21207). Cells were washed with PBS and stained with 1:1000 DAPI for 15 minutes. Images were collected using a Molecular Devices ImageXpress.

#### Proximity ligation assay

Proximity ligation assay was performed using the NaveniFlex Cell Red kit (Navinci, 60025) according to manufacturer instructions with all volumes following recommendations for a 96-well plate. Untreated, scramble shRNA, and SP1 shRNA-treated iPSC-aCMs were fixed in 4% PFA for 20 minutes and washed with PBS. Cells were permeabilized in 0.2% TritonX-100 for 15 minutes at room temperature, then incubated with Naveni Block for 60 minutes at 37°C.

Mouse monoclonal NR2F2 (R&D Systems, H7147) and rabbit monoclonal SP1 (Cell Signaling Technology, 9389) were diluted 1:400 in Naveni Dilutent, and cells were incubated in primary antibody solution for 60 minutes at 37°C. Cells were washed with TBS-T three times. Cells were incubated in 1:40 Navenibody M1 and 1:40 Navenibody R2 for 60 minutes at 37°C then washed with warmed TBS-T three times. Cells were incubated in 1:40 Naveni Enzyme 1 for 60 minutes at 37°C, washed three times with TBS-T, incubated with 1:40 Naveni Enzyme 2 for 90 minutes at 37°C, and then washed with TBS three times. Finally, cells were incubated with 1:1000 DAPI for 15 minutes. After imaging, cells were counterstained with cTnT, ANP, and MLC2V according to the immunofluorescence protocol.

### Image analysis

Single-cell image segmentation and analysis was performed using CellProfiler^40^. First, nuclei were identified from DAPI staining. Cardiomyocytes were then identified by propagating from nuclei to the border of cTnT staining and thresholding based on size based on a previously validated pipeline^41,42^. Measurements of integrated intensity per cardiomyocyte were then collected for cTnT, ANP, and MLC2V. We compared the mean intensity per cells across wells treated with scramble, NR2F2, and SP1 shRNA. We also collected per-well summaries by calculating the median within a well of the per-cell median intensities.

For the quantification of PLA signal, images were segmented using Cellpose (4.2.1.1), with nuclei being labeled using DAPI staining and cytoplasm being labeled using cTnT staining. Nuclei segmentation masks were then imported into CellProfiler. Cells were filtered to cardiomyocytes based on cTnT signal. Puncta were labeled based on intensity and size of signal from PLA. The puncta were mapped to nuclei, enabling single-cell correlations with atrial marker ANP.

## Results

### Human fetal heart gene regulatory networks predict SP1 as a driver of atrial identity

To constrain the architecture of inferred gene regulatory networks (GRNs), we first used CellOracle^20^ to define base GRNs constrained by either TF binding motifs within annotated promoters or within open chromatin, based on human fetal heart scATAC-seq^29^ of atrial and ventricular cardiomyocytes (each pooled across post-conception weeks 6, 8, 12, and 19). We then applied network edge inference based on human fetal heart scRNA-seq^4,18,27,28^ from atrial or ventricular cardiomyocytes (each pooled across post-conception weeks 6, 8, 12, and 19), resulting in both promoter-informed and ATAC-informed versions of GRNs for atrial cardiomyocytes (aCMs) and ventricular cardiomyocytes (vCMs) (**Figure 1A**). Directed edges from TFs to target genes are thus based on a TF-gene expression regression constrained by TF binding motifs within either accessible chromatin or annotated promoter (see **Methods**).

**Figure 1.**
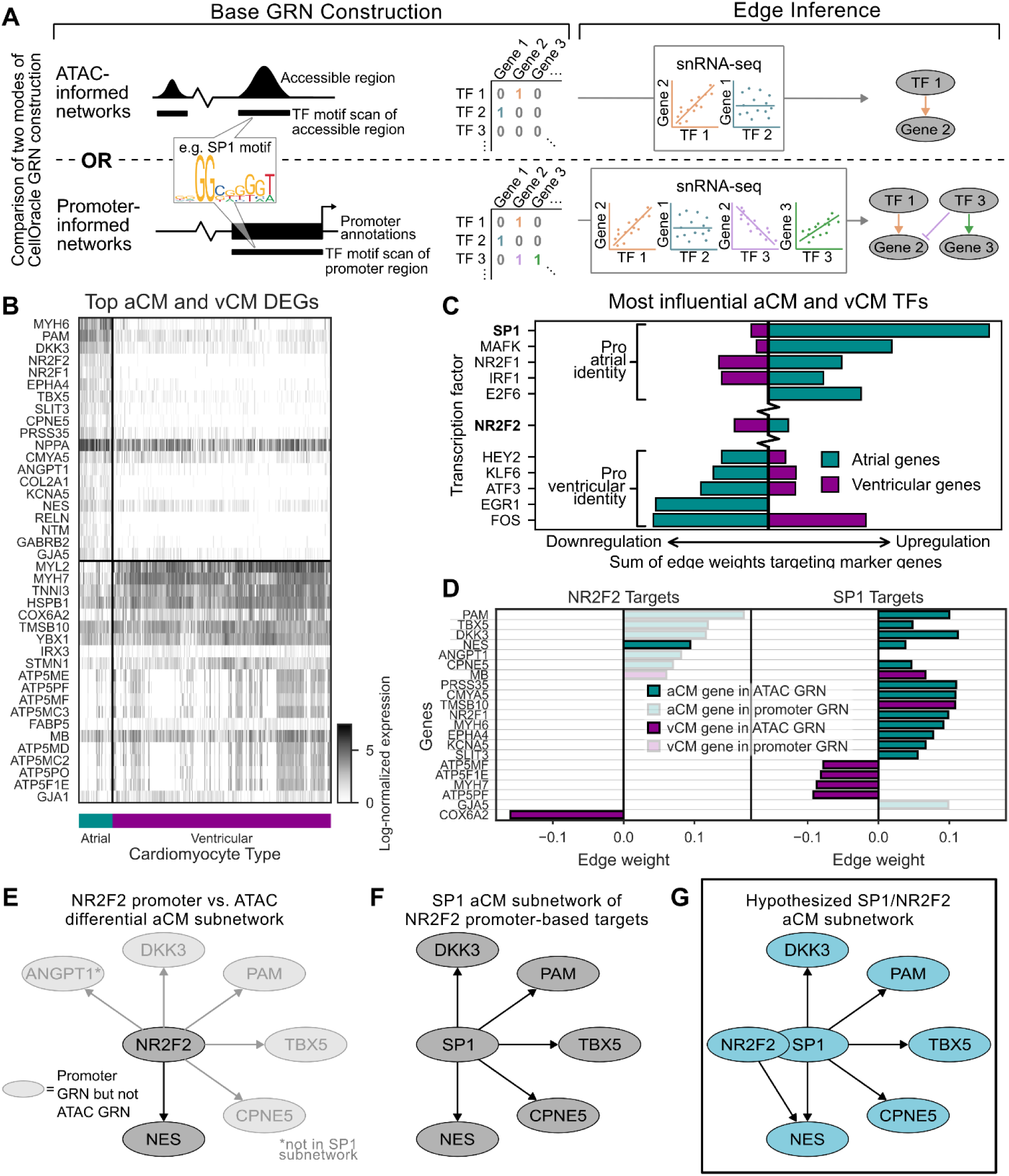
Comparison of human fetal cardiac gene regulatory networks predicts that NR2F2 regulates atrial identity through interaction with SP1. (A) Gene regulatory networks (GRN) were constructed from publicly available human fetal atrial and ventricular cardiomyocyte single-cell RNA sequencing data, with TF-gene edges constrained by either ATAC-based accessible regions of DNA or annotated promoter regions.(B) Top 20 most differentially expressed genes in atrial vs. ventricular cardiomyocytes, from scRNA-seq of human fetal heart.^4,27^ (C) The ATAC-based atrial GRN was used to rank TFs based on their predicted effects on atrial vs. ventricular gene expression, quantified as the sum of edge weights connecting each TF to the top 20 atrial and top 20 ventricular DEGs. NR2F2 shown for comparison. (D) Weight of edges of SP1 targeting top atrial and ventricular DEGs in atrial ATAC-informed or promoter-informed networks. (E) 6 of the 7 genes NR2F2 targets in the promoter-based atrial network are not NR2F2 targets in the ATAC-based atrial network. (F) SP1 targets 5 of the genes that NR2F2 targets in the promoter-based atrial GRN only. (G) Hypothesized cooperation between SP1 and NR2F2 required for atrial identity.

To identify genes that distinguish human fetal atrial vs. ventricular cardiomyocyte identity, we performed differential analysis of the pseudo-bulked scRNA-seq (**Figure 1B**). Many of the top 20 most differentially expressed genes were previously shown to have higher relative expression in aCMs (e.g. MYH6, NR2F2) vs. cVMs (e.g. MYL2, MYH7).

We next sought to identify TFs that are most influential in driving atrial vs. ventricular identity. Examining the ATAC-informed aCM GRN, we computed an atrial-ventricular net identity score that ranks TFs based on the sum of edge weights from that TF to the top 20 atrial and ventricular target genes from **Figure 1B**. This analysis predicted SP1 as the top regulator of atrial identity (**Figure 1C**). While SP1 is a ubiquitously expressed TF that regulates gene expression across a wide variety of processes^43^, to our knowledge it has not been previously characterized as a determinant of atrial identity.

### Comparison of gene regulatory networks inferred by two methods predicts NR2F2-SP1 interaction

Despite being the only TF known to be essential and sufficient to drive expression of atrial identity genes,^6,7^ NR2F2 was not identified by this ATAC-informed aCM GRN as a direct regulator of atrial identity in the human fetal heart. Previous studies characterizing NR2F2 as a regulator of cardiomyocyte identity include NR2F2 binding at TBX5 in E13.5 mouse atrial tissue,^6^ NR2F2 binding at KCNA5 in human ES-derived aCMs,^44^ and NR2F2 targeting of ANGPT1 in E8.5 inflow tract.^17^ Thus, NR2F2 has been shown to bind to regulatory regions of multiple atrial genes in developing cardiomyocytes.

The lack of NR2F2 regulation of atrial genes predicted by the ATAC-informed network could indicate that during fetal heart development, either the NR2F2 binding motifs in the regulatory regions of genes are not accessible, or NR2F2 expression is not correlated with the expression of the target genes. To discriminate between these hypotheses, we compared the ATAC-informed GRNs (that require accessible enhancers or promoters) to the promoter-informed GRNs (that only additionally require correlated gene expression). Network inspection revealed that the ATAC-informed network lacked edges from NR2F2 to top atrial genes that were all strongly targeted in the promoter-informed GRN: PAM, TBX5, DKK3, NES, ANGPT1, and CPNE5 (**Figure 1D**). Thus, NR2F2 has binding motifs and correlated gene expression, but it is not directly accessible to regulatory regions of these genes in the human fetal heart. In contrast, the ATAC-informed aCM GRN contained strong edges from SP1 to these atrial genes (**Figure 1D**).

SP1 and NR2F2 are known to cooperate to drive gene expression in various systems outside the heart,^45–50^ indicating potential for such cooperation in atrial identity. Indeed, of the six atrial genes targeted by NR2F2 in the promoter-based GRN (**Figure 1E**), SP1 is predicted to target five of them in the ATAC-based GRN: PAM, TBX5, DKK3, NES, and CPNE5 (**Figure 1D and 1F**). Therefore, we hypothesized that NR2F2 regulates downstream gene expression of atrial genes through cooperation with SP1 (**Figure 1G**). We predict that this interaction is required for atrial identity during human fetal development.

### SP1 is required for atrial identity

Next, we sought to experimentally validate the GRN-driven hypothesis that SP1 is required for atrial gene expression. We differentiated human iPSCs toward a mixed population of atrial and ventricular cardiomyocytes and knocked down SP1 during the establishment of identity using shRNA lentiviral vectors. We then measured the effect of SP1 knockdown on atrial marker ANP and ventricular marker MLC2V using immunofluorescence (**Figure 2A**). SP1 knockdown resulted in a significant reduction per-well median ANP expression (p=0.0056) but not a significant change in per-well median MLC2V expression (p=0.41) (**Figure 2B and C**). Furthermore, the decrease in ANP expression was quantitatively substantial in the entire population of atrial cardiomyocytes (**Figure 2D**). Thus, SP1 is necessary for atrial identity in human iPSC-derived cardiomyocytes.

**Figure 2.**
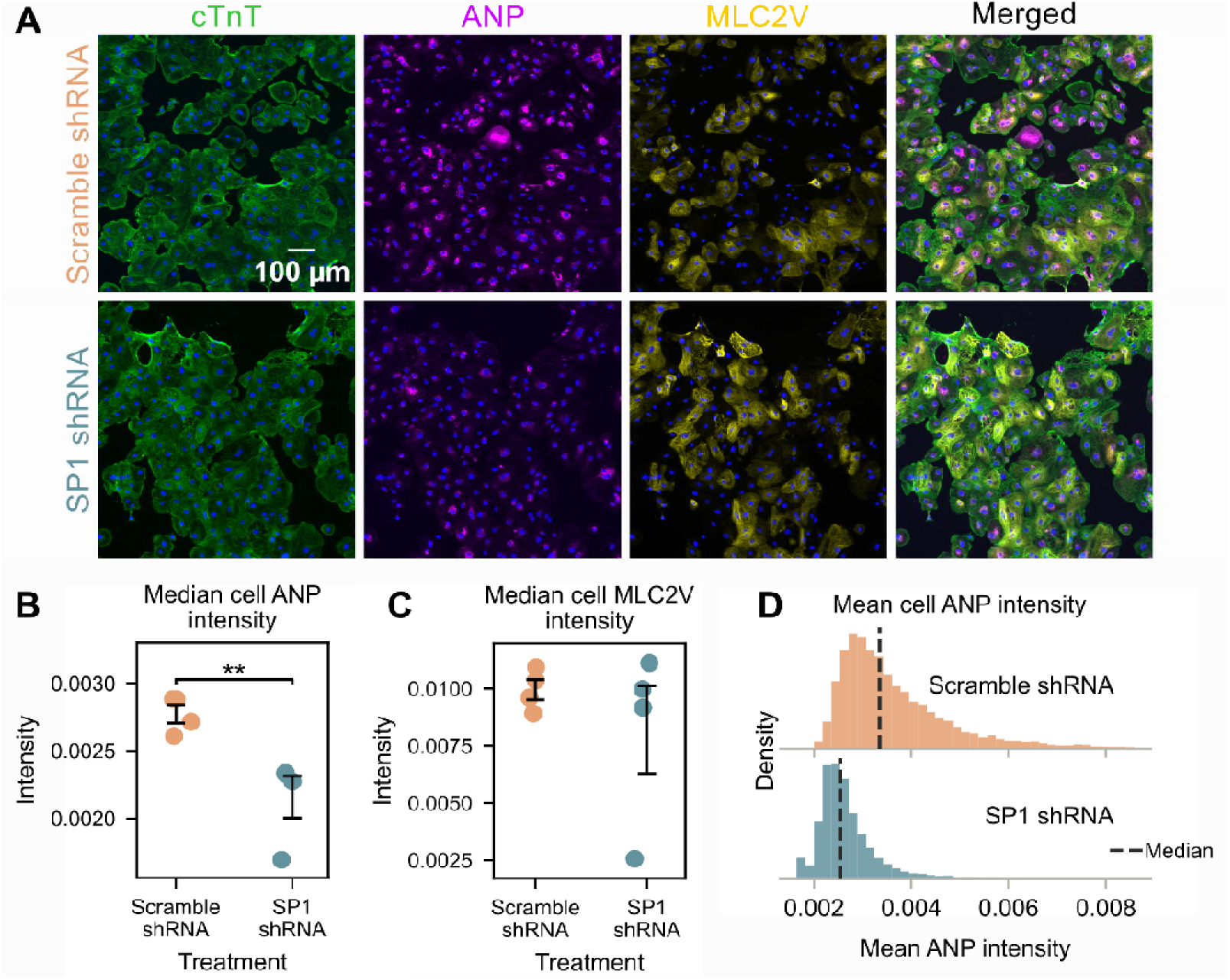
SP1 knockdown reduces atrial identity of human cardiomyocytes. (A) Human induced pluripotent stem cells (hiPSC) were differentiated toward a mixed population of atrial (aCM) and ventricular (vCM) cardiomyocytes for 20 d. The effects of scramble shRNA and SP1 shRNA on general CM marker cTnT, aCM marker ANP, and vCM marker MLC2V were measured by immunofluorescent staining. Median cell (B) ANP and (C) MLC2V immunofluorescent intensity, with dots representing individual wells. Bars represent mean ± SEM. Statistically significant differences were determined by one-tailed t-test, **p < 0.01. (D) Distributions of mean ANP intensity per cell per shRNA condition.

### SP1 and NR2F2 physically interact at atrial promoters

We next asked whether the functional relationship of SP1, NR2F2, and target atrial genes was supported by protein structure and biophysics. We used AlphaFold to predict the protein structures and interactions of SP1, NR2F2, and 50 bp double stranded DNA sequences containing an SP1 binding motif at promoters of atrial genes where we predict SP1 and NR2F2 to interact (PAM, TBX5, DKK3, NES, and CPNE5). Alone, NR2F2 was predicted to not bind the promoter for CPNE5 (**Figure 3A**) or the other four genes. In contrast, SP1 alone bound to the promoters of all five atrial genes, as did the complex of NR2F2 and SP1. These predicted TF-TF-promoter and TF-promoter interactions were quantitatively supported by AlphaFold’s chain pair interface predicted template modeling (ipTM) score, with scores >0.8 indicating high confidence.

**Figure 3.**
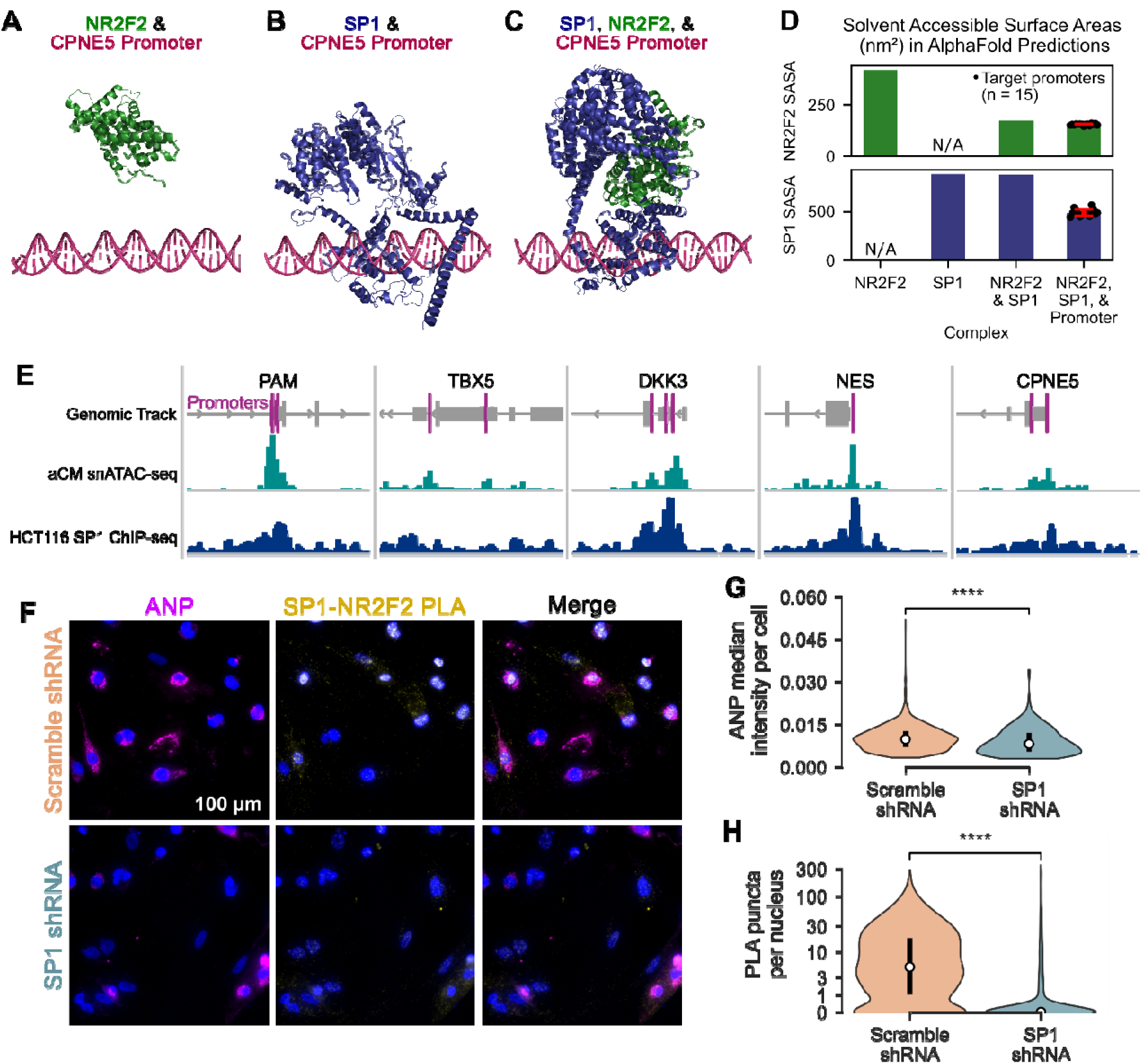
SP1 and NR2F2 molecularly interact at atrial promoters. AlphaFold3 predictions of a CPNE5 promoter (CTCAGGCTGCTCCATCGCCCACCGCACCCCCCACCCCAAATTAGTCAATC) with (A) NR2F2, (B) SP1, and (C) both SP1 and NR2F2. (D) Solvent accessible surface areas (SASA) of NR2F2 and SP1 molecules in AlphaFold-predicted complexes. Lower SASAs indicate greater stability, (E) ChIP-seq from cancer cell line compared with atrial cardiomyocyte snATAC-seq at the promoters of atrial target genes. (F) Proximity ligation assay of NR2F2 and SP1 in human iPSC-derived atrial and ventricular cardiomyocytes, also stained for ANP. (G) Distribution of ANP median intensity per cell. (H) Number of PLA puncta per cardiomyocyte nucleus.

Functional protein interactions are expected to compact protein structure, resulting in a lower solvent accessible surface area (SASA). Consistent with this expectation, SASA for NR2F2 decreased when in complex with SP1 and did not decrease further when in complex with any of the 15 promoters from 5 atrial genes (**Figure 3D**). In contrast, binding to NR2F2 alone did not affect the SASA of SP1, which was decreased only when in complex with both SP1 and promoter DNA (**Figure 3D**). AlphaFold predicted that SP1 bound DNA at its three zinc finger motifs.

To experimentally validate that SP1 indeed binds at the accessible regions of promoters of atrial genes, we compared publicly available SP1 ChIP-seq from HCT116 cells to the aCM scATAC-seq at PAM, TBX5, DKK3, NES, and CPNE5 promoters. At the accessible regions of four of the five the promoters, SP1 had significant binding (**Figure 3E**).

To experimentally validate the predicted physical interaction of SP1 and NR2F2 associated with atrial identity, we performed proximity ligation assays (PLA) between SP1 and NR2F2 antibodies in the mixed human iPSC-derived atrial and ventricular cardiomyocytes. Indeed, we detected PLA signal indicating physical NR2F2-SP1 association specifically in nuclei (**Figure 3F**). PLA signal was reduced when cells were treated with SP1 shRNA, as was ANP signal (**Figure 3G and 3H**).

## Discussion

This study sought to generate mechanistic GRNs of the human fetal heart, providing new insights into drivers of atrial identity. Comparison of GRNs with and without data on single-cell chromatin accessibility led to the hypothesis that rather than acting directly, the canonical atrial transcription factor NR2F2 regulates atrial identity indirectly by forming a transcriptional complex with SP1. We demonstrate that during human heart development, SP1 is accessible to gene regulatory regions and correlates with expression of genes that define atrial identity. Network predictions of an NR2F2-SP1 transcriptional complex that drives atrial identity were validated through structural predictions in AlphaFold3 and in human iPSC-derived atrial cardiomyocytes via effects of SP1 knockdown and physical association of NR2F2 and SP1 by proximity ligation. These analyses and validation experiments support SP1 and its tethering to NR2F2 as a driver of atrial cardiomyocyte identity.

NR2F2 is the most well-characterized TF for specifying atrial identity. It binds to selected atrial genes in early developing mouse heart, and its knockdown shifts atrial cardiomyocytes towards ventricular identity.^6,15,44,51^ Therefore, the unavailability of open chromatin for direct binding of NR2F2 at atrial genes in data from human fetal hearts at 6 to 19 weeks post-conception was unexpected. SP1 and NR2F2 have been shown to cooperate to drive downstream gene expression in various cell types and species.^6,45–50^ For instance, NR2F2 tethering to SP1 regulates E2F1 expression in HUVEC cells and NGFI-A expression in rat urogenital mesenchymal cells.^46^ In E13.5 mouse cardiomyocytes, previous work demonstrated that deletion of an Sp1 motif at the Tbx5 locus reduced its expression, suggesting that Nr2f2 regulates Tbx5 expression using the site of the Sp1 binding motif^6^. However, to our knowledge SP1 and NR2F2-SP1 tethering have not previously been identified as necessary for atrial vs. ventricular identity.

While the ATAC-informed GRN showed no NR2F2 binding motifs in the accessible regions of the atrial target genes during this stage of development, those promoters contain NR2F2 binding motifs. NR2F2 may directly bind to those sites at earlier or later stages of development, potentially helping to recruit SP1 to those sites.^6,46^ Analysis of binding at earlier timepoints would be required to determine the dynamics at the initiation of NR2F2-SP1 tethering.

Using sequence-to-structure prediction with AlphaFold, we predicted that SP1 binds to atrial gene promoters and that NR2F2 binds SP1 to form a compact ternary complex. This biophysical prediction helps validate the GRN predictions. Further, AlphaFold-generated interactions predict that NR2F2 does not independently bind these sites, making SP1 necessary for NR2F2 to regulate gene expression via these sequences. PLA signal for NR2F2-SP1 interaction correlated with ANP at the single-cell level. Thus these findings predicted and experimentally validated NR2F2-SP1 tethering as important for aCM identity. Some non-CM (cTnT negative) cells had nuclear PLA puncta as well, indicating that SP1 and NR2F2 interact in other cell types consistent with previous reports of this complex in microglial, mesenchymal, and endothelial cells^45–47^. Thus, while NR2F2-SP1 tethering is not unique to cardiomyocytes, we find that within developing cardiomyocytes this complex enables SP1 to be a strong driver of atrial vs. ventricular identity.

This study develops a novel method of comparing GRNs inferred using context-specific snATAC-seq with a context-independent promoter-based approach. We demonstrate that this method reveals that the canonical atrial TF NR2F2 does not directly bind to atrial promoters during late morphogenesis and early maturation of the heart, instead driving atrial identity by cooperating with SP1. This network comparison reveals that use of ATAC-informed GRN alone may fail to capture regulatory interactions between TFs such as TF-TF tethering. Gene regulation is highly complex and concerted, and we are just beginning to map the cooperative effects of TFs. The approach of comparing ATAC-informed and promoter-informed GRNs may be generalized to systematically identify other cooperative TFs driving gene expression.

## Funding

This study was funded by the National Institutes of Health grants HL176700, HL160665, HL163925 (to JJS), T32HL007284 (to APC), a University of Virginia Harrison Undergraduate Research Award (to PNG), Coulter Foundation Postdoctoral Research Grant (to APC), and American Heart Association 25POST1356494 (to APC).

## Notes

### Competing Interest Statement

The authors have declared no competing interest.

